# Two notorious nodes: a critical examination of MCMCTree relaxed molecular clock estimates of the bilaterian animals and placental mammals

**DOI:** 10.1101/2022.07.01.498494

**Authors:** Graham E. Budd, Richard P. Mann

## Abstract

The popularity of MCMCTree for Bayesian inference of clade origin timings has generated several recent publications with focal results considerably older than the fossils of the clades in question. Here we critically examine two such clades; the animals (with focus on the bilaterians) and the mammals (with focus on the placentals). Each example displays a set of characteristic pathologies which, although much commented on, are rarely corrected for. We conclude that in neither case does the molecular clock analysis provide any evidence for an origin of the clade deeper than what the fossil record might suggest. In addition, both these clades have other features (including, in the case of the placental mammals, proximity to a large mass extinction) that allow us to generate precise expectations of the timings of their origins. Thus, in these instances the fossil record can provide a powerful test of molecular clock methodology, and why it goes astray.

## Introduction

The apparent clash between molecular clock estimates of the time of origins of clades versus what the fossil record might be thought to say has been a significant source of anxiety in the last two decades^1–4^. One of the reasons for this, perhaps, is that many molecular clock practitioners come from a perspective where the fossil record may seem to be a haphazard and unreliable source of information about when clades arose. After all, the discovery of *Latimeria* c. 70 million years after the youngest coelacanth fossil^5,6^ is a salutary reminder that the fossil record can sometimes have very significant lacunae. Nevertheless, the fossil record clearly shows significant structure that would be difficult to reconcile with such *general* unreliability^7,8^. Biostratigraphic division of rocks on the basis of biotic succession is sometimes possible to a precision of a few hundred thousand years or less. For example, a recent study of ancestor-descendent relationships within graptolites suggested that the modal gap between the last appearance of an ancestral graptolite and the first appearance of its descendent was as little as 49 kyrs^9^. Given that some of this time will be taken up by non-appearance of the ancestor, it follows that the descendants appear in the fossil record only a few tens of thousands of years after their true emergence, which on most geological time scales is essentially simultaneously. In addition, our ability to map rapid post-extinction radiations^10^ such as at least ordinal level placental mammals strongly suggests that in some cases, the fossil record must be giving us a more or less accurate picture of temporal evolution.

Molecular clock estimates, which have been argued to have essentially taken over from the fossil record as the ultimate measures of evolutionary time^11^, conversely, often give much larger gaps between first appearances of clades and the fossil record. To give some recent examples of the estimated gaps between the first recognised fossil of a clade and the 95% credible interval of molecular clock estimates: beetles (55–134 Myrs older than the oldest fossil)^12^; sand dollar and sea biscuit echinoids (40–65 Myrs)^13^; cheilostome bryozoans (132– 238 Myrs)^14^; angiosperms (23–121 Myrs)^15^ and bilaterians (50–143 Myrs)^16^. The possible reasons for these sorts of large gaps have been much discussed, but here we wish to focus on two of them: the way that calibration ranges are converted to priors, and the influence that priors have on the outcome. Whilst these issues have also been the subject of considerable discussion in the literature^17,18^, we do not believe that these discussions have been followed through to their logical conclusion. As a focal point for our investigation, we take two rather notorious nodes; the origin of the animals^16^, and the place of the bilaterians within them, and the origin of the placental mammals^19^; both of which have generated considerable controversy. There are of course several different types of molecular clock methodology, some of which we have critiqued^20^ earlier; here we focus on MCMTree analyses in the PAML package^21^ and the conditional prior method^22^ of incorporating fossil dating evidence into trees. This sort of analysis has been the source of much proposed revision to our beliefs about the origins of various clades, so is a worthwhile topic for a more detailed discussion.

### Animal origins

The most complete examination of the origin of animals from an MCMCTree perspective remains the comprehensive study of Dos Reis et al. (2015)^16^. This study used four different calibration strategies to generate priors for analyses that were run under autocorrelated and uncorrelated rate conditions, and with a variety of rate partitions in the data. Under this wide range of conditions, they recovered dates for the origin of the animals between 833 and 650 millions years ago, and bilaterians between 688 and 596 million years ago. In contrast, the oldest generally accepted animal fossils are younger than about 575 Ma^23^. The commencement of the fossil record of the bilaterians has some uncertainty in it, because there is no consensus about basal bilaterian morphology; but it is widely accepted that trace fossils younger than 560 Ma represent at least total-group bilaterians^20,24,25^; and that body (and small carbonaceous) fossils close to the base of the Cambrian represent bilaterian clades such as total-group chaetognaths (i.e. protoconodonts) or scalidophorans^26,27^. It has also been argued that the trace fossil *T. pedum*, which marks the base of the Cambrian, but drifts below it, was made by a priapulid-like animal^28,29^. A fossil-based date of c. 555-540 Ma for the origin of the crown-group bilaterians would thus not be unreasonable. In both cases, therefore, a considerable gap exists between the youngest molecular clock estimates and the fossil record. The calibration strategies that Dos Reis et al. (2015)^16^ used were: a uniform calibration, from the age of the oldest fossil back to a date where no fossil is found even though it reasonably could be; a skew-normal distribution that places the weight of probability close to the oldest fossil; then two Cauchy distributions, one “optimistic” about the fossil record, and one “pessimistic”. For clarity of illustration, we will examine the uniform distributions here. We note, however, that the basal node of the metazoans as a whole was in each of these four cases calibrated with uniform distributions. Uniform calibration ranges are in addition extended by “soft” margins at both ends in order to account for mistakes in fossil assignment or oldest age estimation^30^.

As noted by Dos Reis et al. (2015)^16^, the logical necessity of clades being younger than the clades they are nested within means that initial calibration ranges must be truncated to form a joint prior structure. In the case of the basal nodes of the animals, this truncation is striking (Fig. 1).

**Fig. 1.**
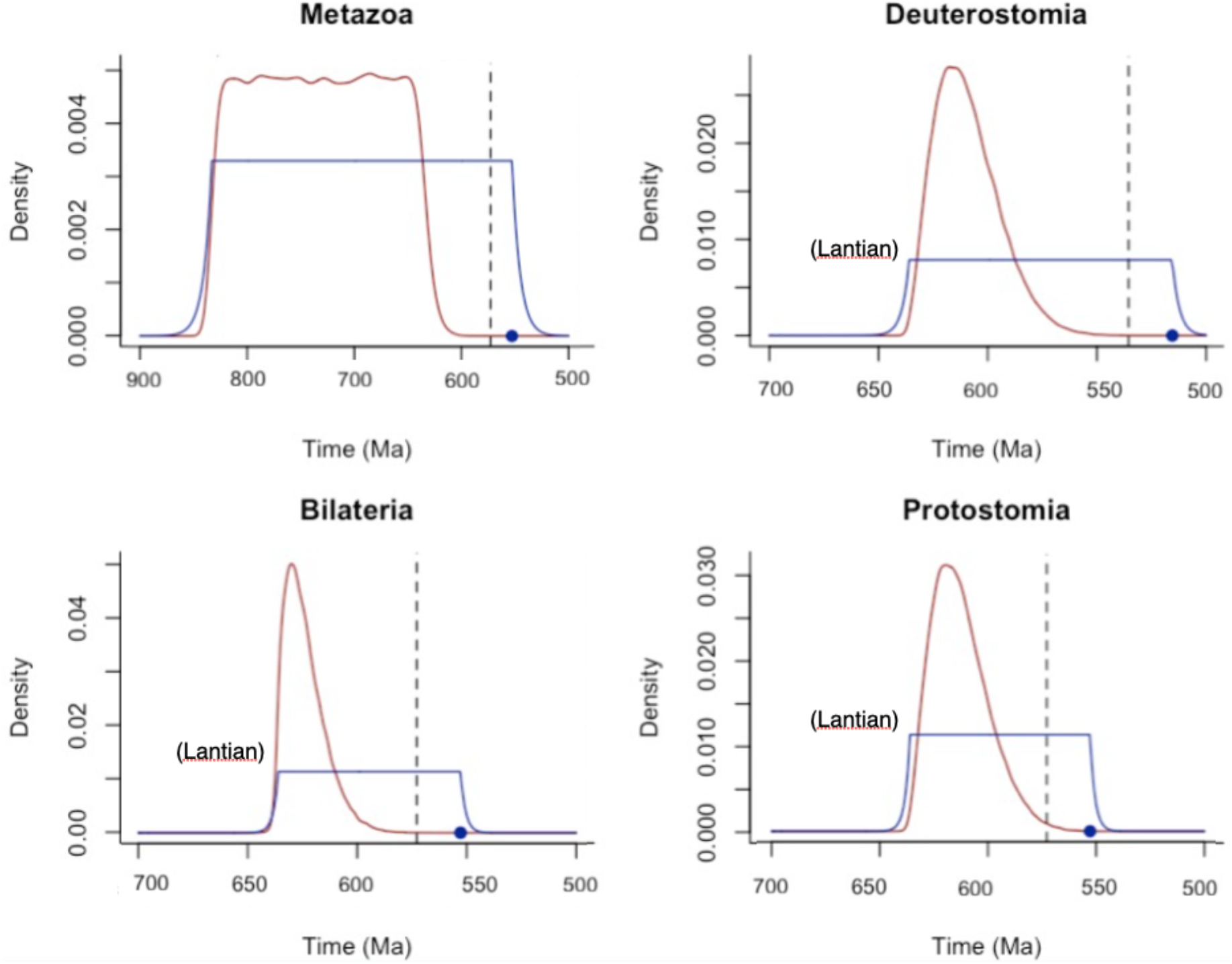
The uniform calibration ranges (blue) and subsequent priors (red) used in Dos Reis et al. (2015)^16^. The oldest fossil used for setting the youngest soft limit is marked with a blue dot; the dashed line is 20 Myrs older than this fossil in each case. Note that the Lantian biota, used as a soft maximum age for three of these nodes, has recently been dated to be c. 602 +/- 8 Ma^31^. Note that in each case, the prior assigns little probability to dates near the oldest fossil.

As can be seen, in each of the Metazoa, Bilateria, Protostomia and Deuterostomia, the prior has a much older, and, perhaps more importantly, narrower age range than the calibration range. Indeed, in all cases apart from the Protostomia, the prior assigns less than 5% probability of an age within 20 Myrs of the oldest fossil. A few notes on the dates used are in order: the soft maximum age used for the Metazoa was the maximum age of the Bitter Springs exceptionally-preserved biota at c. 833 Ma^32^; and the soft maximum for the other three nodes was the Lantian biota at c. 635 Ma. Since Dos Reis et al. (2015) was published, the Lantian has been redated to be younger than c. 602+/- 8 Myrs^31^. In addition, we do not accept that *Kimberella*, used as the oldest protostome, is in fact a protostome^33^. Conversely, we would accept the oldest bilaterian-style trace fossils as the oldest possible crown-group bilaterians at c. 560 Ma^20^. For the sake of simplicity again, we are choosing to bracket possible dates from other putative early metazoans such as the 890 Ma sponge described by Turner^34^, although we do not find these compelling.

The fossil record of the earliest animals seems to show a fairly convincing phylogenetic “unfolding”, starting with total-group animals (or possibly total-group eumetazoans)^35^ in the oldest Ediacaran assemblages at around 575 Ma^36^; then total group bilaterians and crown group bilaterians^10,20,37,38^. Thus, the fossil record, on its surface at least, seems to suggest that animals emerged at perhaps around 580 Ma and rapidly diversified after this. However, the joint prior structure of the analyses of dos Reis et al. (2015) assigns an extremely low probability to this scenario. We would thus argue that their analyses do not provide a convincing test of the hypothesis of the fossil record more or less faithfully recording the origin of the animals^39^; instead the prior structure effectively rules out this hypothesis *a priori*.

### Prior-posterior plots: When the priors and posteriors agree

We introduce here a tool for examining the outcomes of molecular clock analyses, the prior-posterior plot. We note that a similar plot was previously used in Warnock et al. (2015)^18^, albeit in a different context and for a different purpose, and similar sorts of approaches for examining individual nodes have also been made elsewhere^40^. Our plots plot the mean and 95% credible interval for the prior and posterior for each node in an analysis. In the plot of the analysis of the focal result (i.e. with independent rates) of dos Reis et al. (2015) for uniform priors (Fig. 2A), it can be seen that most nodes lie close to or even on the line corresponding to a 1:1 match between the prior and posterior, and have similar credible intervals. In addition, those few nodes that lie far from this line typically have extremely broad posterior ranges, ranging over hundreds of millions of years. This analysis suggests that the *molecular* part of the analysis tells us very little about the age of emergence of the clades in question: posteriors are either uninformatively broad, or closely match the means of the prior.

**Fig. 2.**
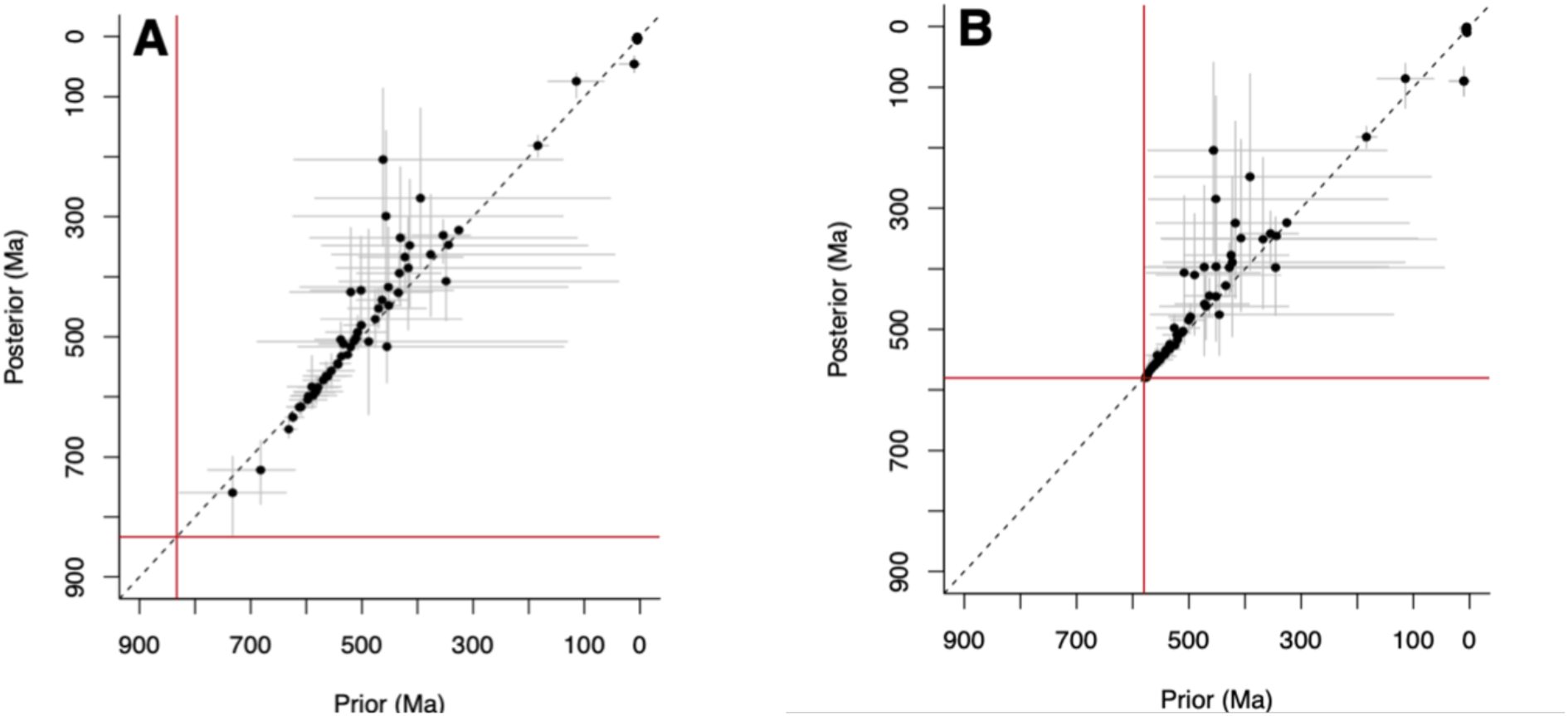
Prior-posterior plots for the analysis of Dos Reis et al. (2015) on the origin of the animals, using uniform calibration ranges with independent rates. Each point represents a node in the phylogeny with it associated 95% HDP range. The red lines in case represent the soft maximum age for the Metazoa (i.e. the root of the tree). **A** The focal analysis of Dos Reis et al. (2015) setting the soft maximum age at c. 833 Ma. **B** A reanalysis setting the soft maximum age at 580, i.e. just before the oldest Ediacaran assemblages^36^.

We then re-ran the focal analysis of dos Reis et al. (2015) but changing a single number, the soft maximum age of the Metazoa, to reflect the diversification seen in the fossil record, i.e. to 580 Ma (Fig. 2B; c.f. ref.^20^). As can be seen, the analysis accommodates this shift simply by bunching up the earliest nodes, including the bilaterians. Once again, there is a close prior-posterior correspondence.

What can be concluded from these plots? The first, and perhaps most troubling, is that the Bayesian inference overall adds little to the probabilities of the priors. In addition, the fact that the priors are algorithmically generated from the calibration ranges and have no biological input into them suggests that the posteriors do not properly reflect our true prior beliefs based on the fossil record. Finally, the flexibility of the rate prior means that the molecular evidence is compatible with the fossil record providing a faithful record of animal diversification. Thus, this analysis does not provide any compelling evidence that either the animals or their subsumed nodes (e.g. Bilateria) are significantly older than their fossil record.

Such an analysis might suggest that the molecular evidence has literally *no* effect on the outcome of the inference. However, the prior rate model does create structure that does eventually constrain the posteriors. For example, if the soft maximum limit for the metazoans is set to 1000 Ma rather than 833 Ma, the mean age for the origin of the metazoans is essentially unchanged (mean age c. 770 Ma vs 760 Ma; data not shown). However, this restraint seems here to act only in one direction, i.e. in restraining very deep estimates for the basal node. We thus now turn to slightly different, and also instructive, example, where the data clearly do exert an important influence on the posteriors, that of the placental mammals.

### When the priors and posteriors disagree: the case of the placental mammals

The origin of the placental mammals has been examined by, among others, Álvarez-Carretero et al. (2021)^19^. This analysis used a huge phylogenomic dataset and concluded with strong probability that the crown placental mammals emerged c. 83.3-77.6 Ma, i.e. considerably before the K-Pg boundary at c. 66 Ma. This is in stark contrast to analyses such as that of O’Leary et al. (2013)^41^ which concluded that the crown placentals emerged rapidly after the K-Pg boundary.

In the Álvarez-Carretero et al. analysis, a uniform calibration for the placentals from the oldest fossil they selected at 61.66 to 162.5 Ma was used, derived, at least partly, from previous molecular clock results^42^.

Once again, the calibration to prior conversion in this case is problematic (Fig. 3A), with an extreme bias towards older dates of the calibration. Sampling the prior 1 million times generated no dates younger than c. 69 Ma (Fig. 3B). Thus, the prior assigns a vanishingly low probability to the placentals emerging after the K-Pg boundary – i.e. much less than one in a million (c.f. ^39^). Nevertheless, this is after all what the fossil record suggests: there are thousands of stem-group fossils known from before the boundary, and no uncontroversial crown-group forms; yet a large number of unambiguous crown group forms appear soon after the boundary^43–45^. The prior-posterior plot in this case is of particular interest, because the posterior of the placentals and nodes near it plot far from their respective priors (Fig. 4). In this instance, the problem is not that the prior controls the posterior, but rather that the prior is so far from what the data suggests. In other words, the “signal” from the prior and the “signal” from the data are fundamentally different; and combining two very divergent signals in a Bayesian context^46^ should acknowledge the plausible scenario that one or other signal (or both) may be systematically erroneous^47^. In this instance, the prior assigns approximately 1 in 11000 probability (0.009%) to ages younger than 83 Ma. That is, the age eventually presented as the focal estimate for the placental node is one that was considered highly unlikely according to the prior.

**Fig. 3.**
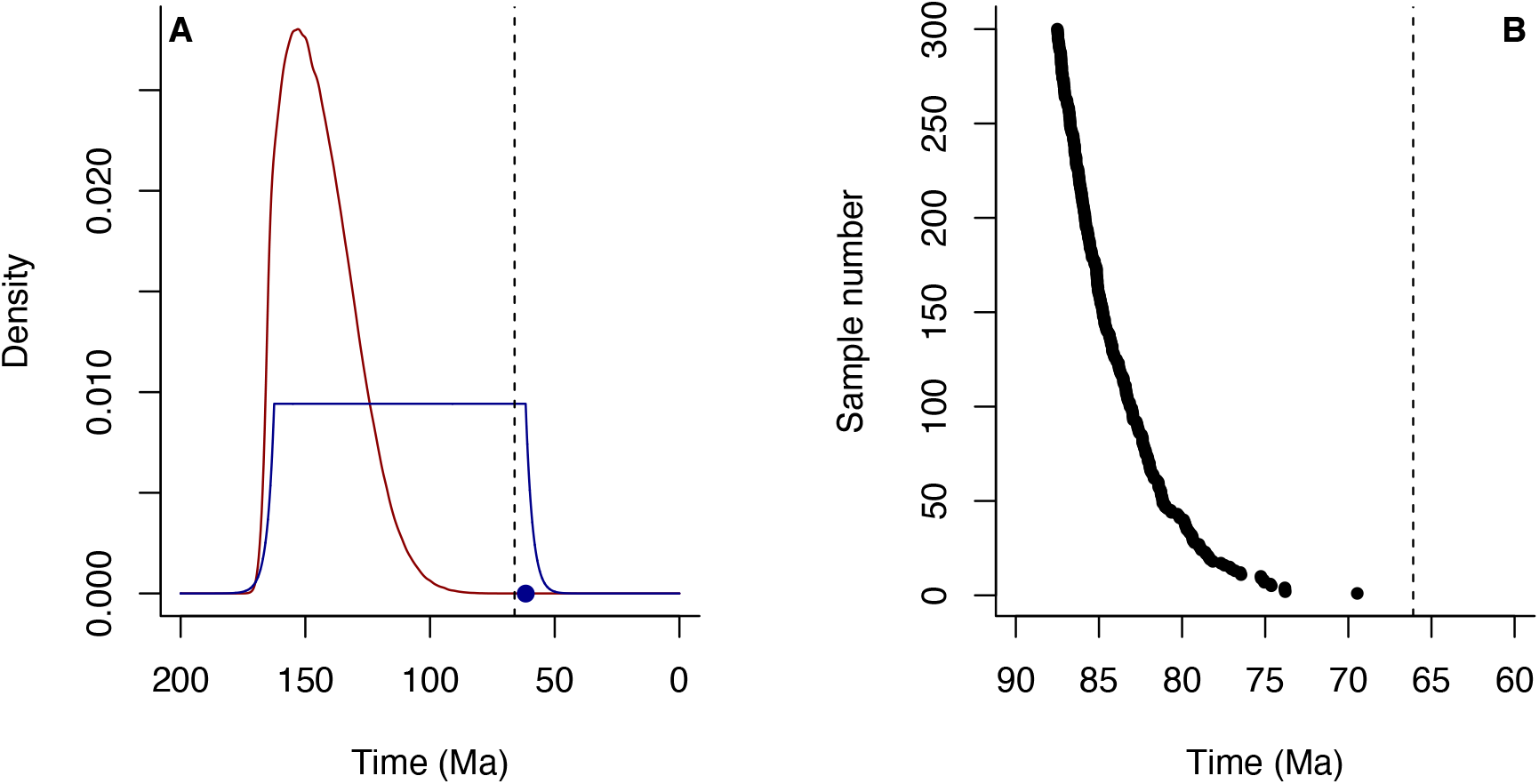
**A** Calibration to prior conversion in the analysis of mammal origins of Álvarez-Carretero et al. (2021). Dashed line: K-Pg boundary; blue: uniform calibration range with soft limits; blue dot, oldest fossil; red: prior. **B** The youngest 300 of one million samples of the prior, showing the vanishingly low probability of an appearance close to or after the K-Pg boundary.

**Fig. 4.**
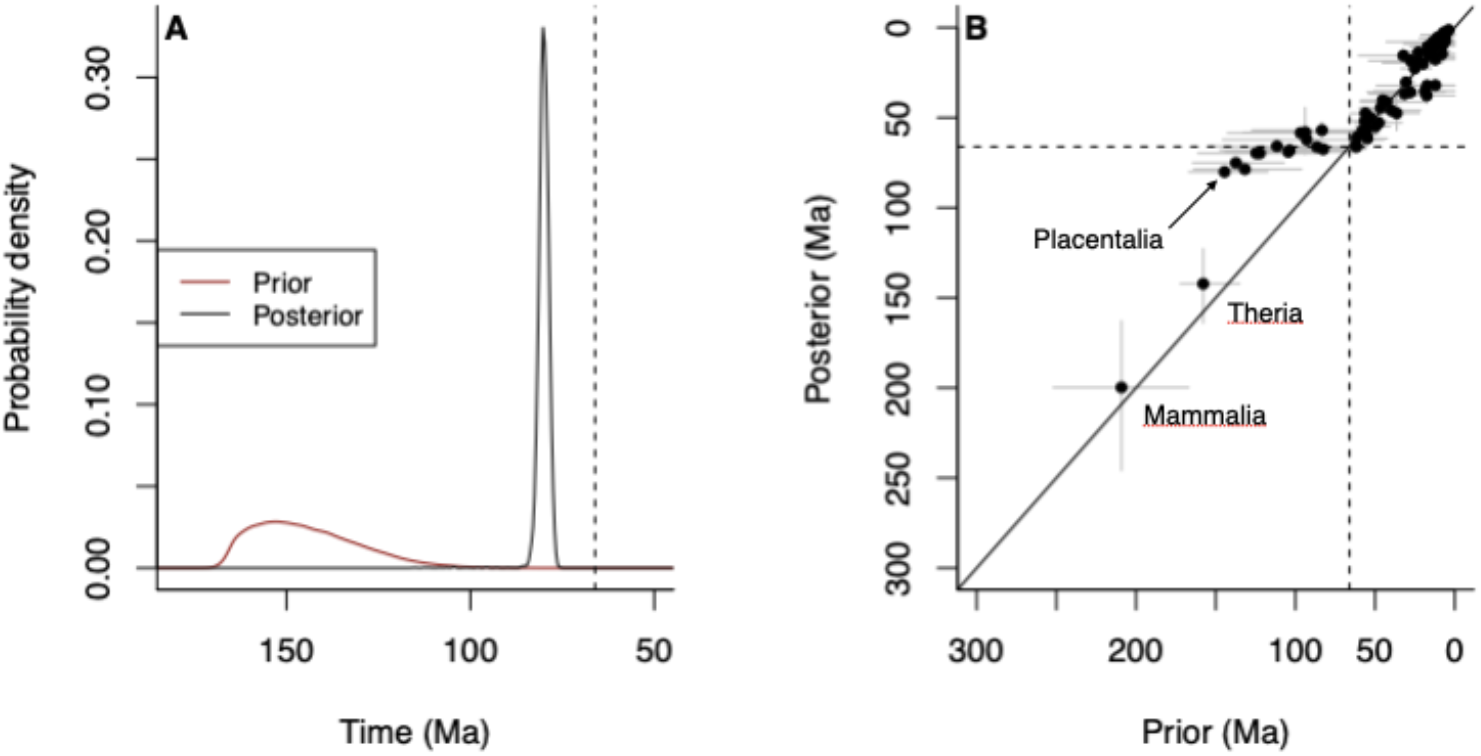
**A** The prior and posterior distributions for the placental mammals from Álvarez-Carretero et al. (2021). Note that in this case, the posterior lies far outside the prior distribution. **B** The prior-posterior plot for the entire analysis (72 taxa). Note that the nodes close to the placentals trend off the prior-posterior equivalence line, a probable result of the very deep placental prior; the large gap between the Theria and the placentals, and the large number of nodes constrained by their priors to lie after the K-Pg boundary.

Álvarez-Carretero et al. (2021) tested the influence of the priors by using two types of truncated Cauchy calibrations (their extended data figure 2), and showed that when a very sharply-defined Cauchy calibration was used for the placentals, the mean age of appearance was only a few million years younger than that of the focal analysis, with the implication that choice of calibration had only limited influence. We consider this result to be problematic, however. Truncated Cauchy distributions have undefined means and variance, as they do not converge, which implies that they are very heavy tailed. Indeed, when comparing the focal prior to their “short” truncated Cauchy prior, one can see that the Cauchy prior starts assigning more probability than the focal prior as deep as c. 93 Ma (Fig. 5). In addition, the gap between the placentals and the next deepest node, the Theria, is considerable, so that the placentals are effectively the root of the clade. In such circumstances, use of a Cauchy prior is hazardous^6^ because it is easy for the posterior to drift deep.

**Fig. 5.**
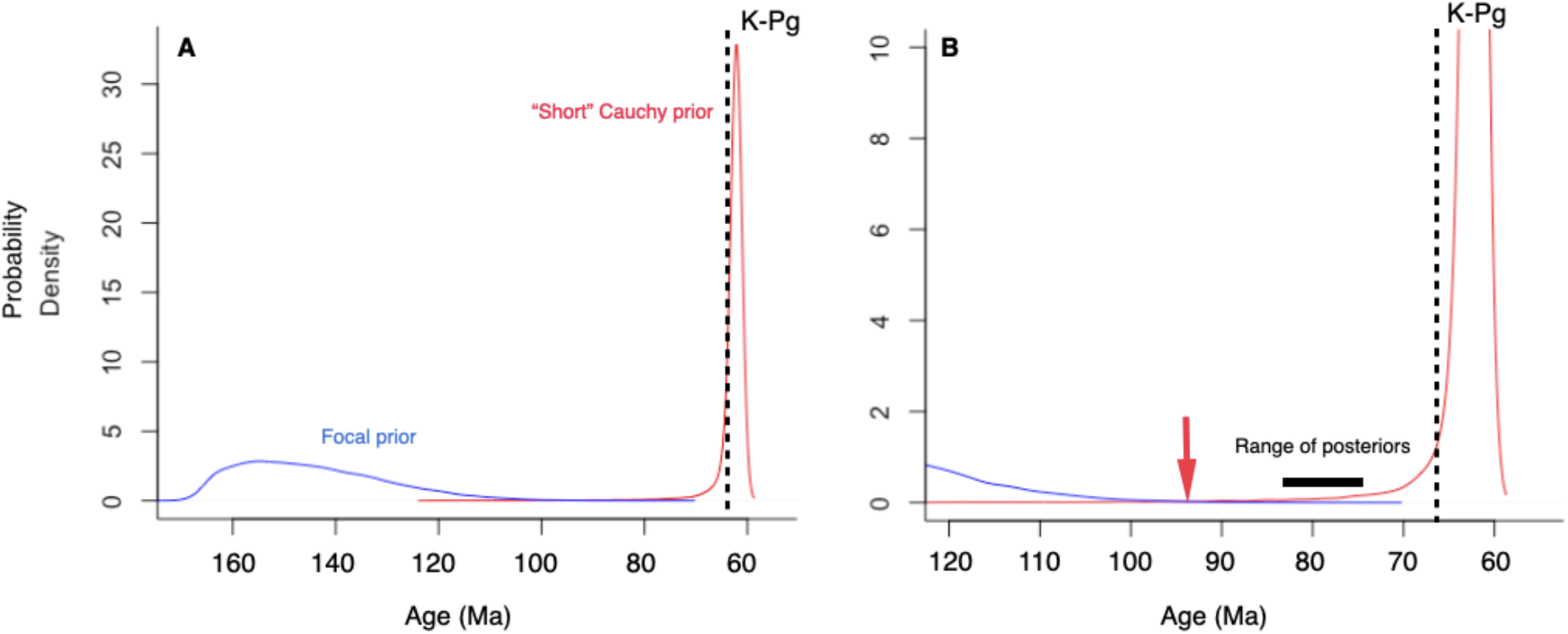
**A** The “short” truncated Cauchy prior of Álvarez-Carretero et al. (2021) (red) plotted together with the prior of the focal analysis (blue). Note that when zoomed in (**B**) the Cauchy prior can be seen to assign more probability than the focal prior for ages as deep as c. 93 Ma (red arrow), older than any of the posteriors.

Rather than change the *type* of prior, then, we re-ran the analysis using a uniform calibration for the placentals and other deep nodes within them from the age of their oldest fossil down to the K-Pg boundary, with soft bounds on both ages (Fig. 6). We note that similar calibration ranges are used for 11 other nodes in their analysis. Once again, construction of the placental prior from this calibration range led to a skewed distribution almost exactly on the boundary (mean c. 65.53 Ma, hdi 64.30–66.44 Ma). Under such conditions, the analysis recovered a posterior mean age very close to the age of the boundary (mean 66.20 Ma, 95% hdi 65.60–66.96 Ma for uncorrelated runs, and mean 66.55 Ma, 95% hdi 65.88–67.59 Ma for correlated runs). In both cases, the posterior is compatible with a post-K-Pg origin of the placentals.

**Fig. 6.**
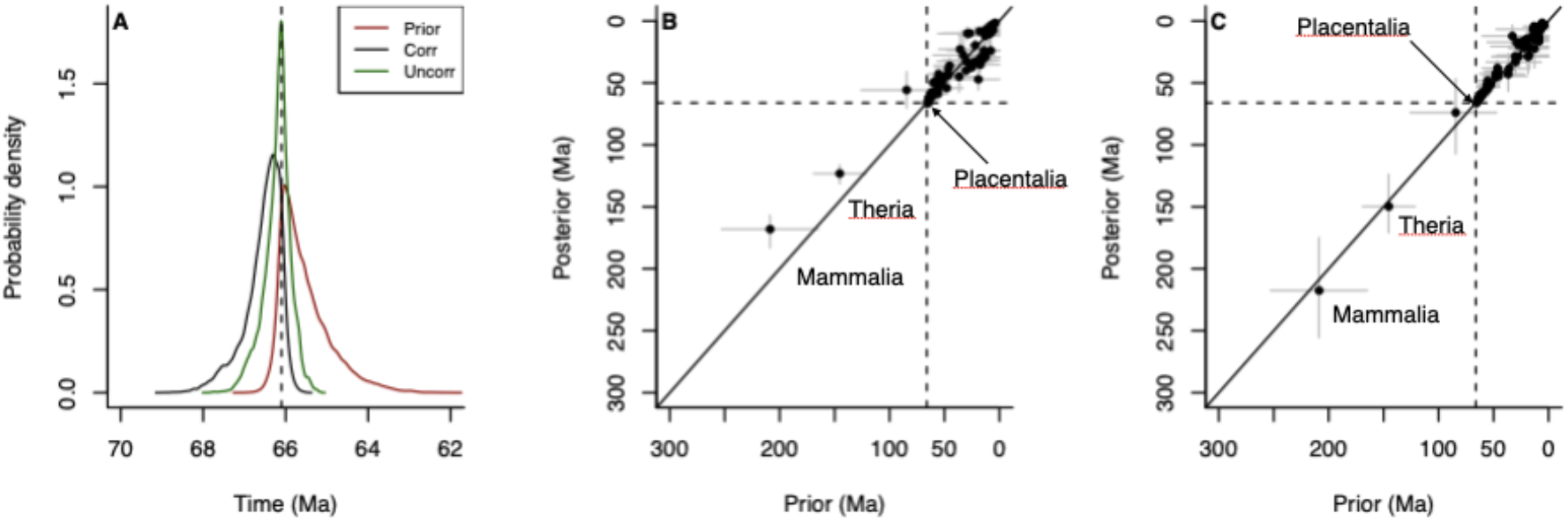
The focal analysis of Álvarez-Carretero et al. (2021), rerun with a uniform calibration on the age of the placentals down to the K-Pg boundary, with soft limits. **A** the prior and the posteriors for uncorrelated and autocorrelated runs for the placental node. **B** prior-posterior plot for the mammals run with an autocorrelated rate model. **C** prior-posterior plot for the mammals run with an uncorrelated log-normal rates model. Note that under such conditions, the posteriors return to plotting broadly along the 1:1 prior-posterior line, and that the molecular evidence provides no compelling evidence for a pre-K-Pg boundary emergence of the placentals.

We note that in the case of the “correlated” run (i.e. with rates on lineages being autocorrelated via geometric Brownian motion (GBM)), both the two old nodes and some of the younger nodes plot further from the prior-posterior line than for the independent (“uncorrelated”) log-normal rates model. Álvarez-Carretero et al. (2021) showed with their priors that the GBM model was more likely than the independent log-normal one. Whilst we have not assessed the marginal likelihood for our priors, we note that mass extinctions (see below) should be expected to create a distinct structure in the spacing of lineages before and after them, as post-extinction species are more likely to lead to present day survivors than pre-extinction ones. This implies that lineages before the mass extinction are likely to represent long periods of time with many implied internal speciation events that gave rise to now extinct plesions, whereas lineages after it may arise more rapidly and may thus more closely approximate to the historical pattern of speciation. Under such circumstances, one would not expect close rate correlation between pre- and post-extinction lineages, as it is presumably the inheritance of rates from ancestral species and not ancestral lineages p*er se* that creates the correlation structure.

The case of the placental mammals is instructive because although their emergence before the K-Pg boundary is widely accepted, there are strong theoretical and empirical grounds for thinking that they really emerged *after* it. The first is provided by the lineage-through-time (LTT) plot (Fig. 7). As can be seen, when the placental emergence is placed where Álvarez-Carretero et al. (2021) suggest it is, the lineages rapidly increase across this boundary with no apparent perturbation. However, it is widely accepted that the mammals were subject to an extremely severe extinction at this point, with numbers up to 95% species extinction being suggested^49^. The slope of the LTT plot is equal to the rate of speciation (λ) multiplied by the probability of survival to the present. Under homogeneous conditions of constant speciation and extinction the probability of survival to the present stays fairly constant until close to the present^8^. However, when these conditions are punctuated by mass extinctions the survival probability is substantially greater after the mass extinction than before, typically leading to an increase in the rate of lineage creation^9^. For the LTT plot to pass unperturbed through the extinction would imply that there must be an immediate and sustained decrease in speciation rate after the mass extinction that exactly balances the increase in survival probability, such that lineages continue to be produced at the same rate. This is inherently implausible and unparsimonious, and the more logical expectation is that the break of slope that is clearly seen in the LTT plot of Álvarez-Carretero et al. (2021) should occur not just before, but *at* the mass extinction^50^. Furthermore, the probability of the crown group forming is, in general, at its lowest just before a mass extinction. This is a direct consequence of the difficulty of coalescence past a mass extinction. Thus although an earlier MCMCTree analysis^53^ reported “uninterrupted” diversification of placental mammals across the boundary, this is both theoretically implausible and contradicted by the fossil record.

**Fig. 7.**
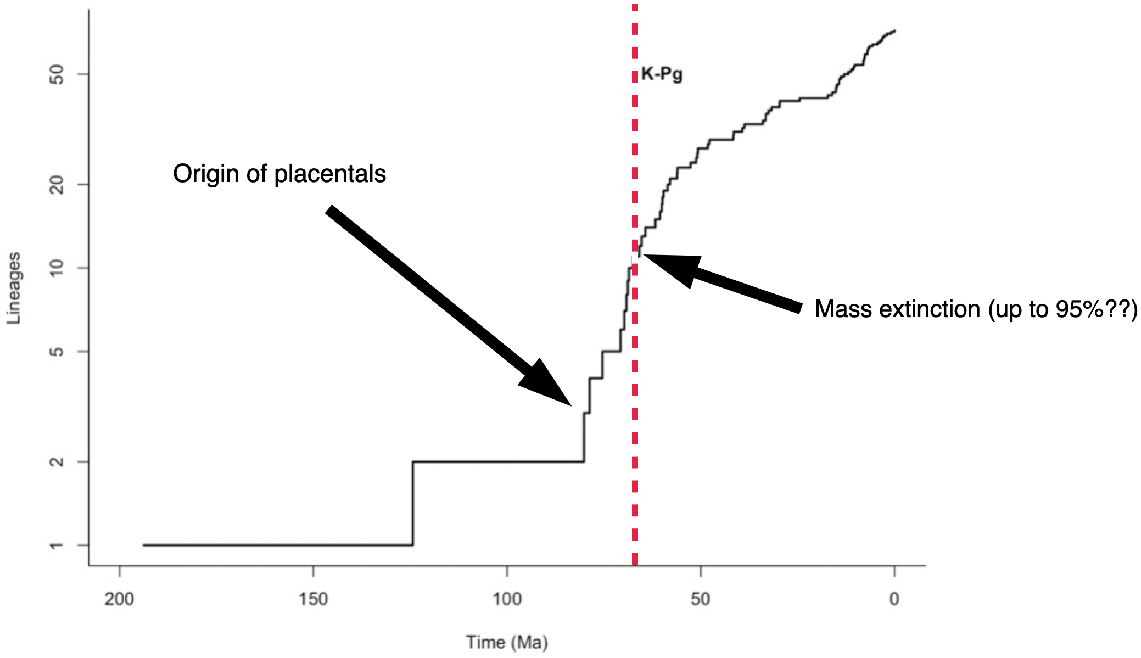
The central lineage-through-time (LTT) plot of Álvarez-Carretero et al. (2021) for 72 taxa. Note that in their dating, the LTT plot passes unperturbed through the K-Pg boundary, despite the very high rates of mammalian extinction at that point.

The emergence of the crown group, and the effect of mass extinctions, can be modelled using birth-death modelling^51,52^. A crown group can logically emerge either before or after a large mass extinction; but it is most likely to do so either early on in the history of the total group, or just after the mass extinction; the *least* likely time of crown-group origin is just before the mass extinction (Fig 8A). What is more, if the crown group forms before the mass extinction we still expect to see a sharp increase in the rate of lineage creation immediately after the mass extinction, creating an inflection point in the LTT plot at 66 Ma (Fig 8 B-C), which is notably absent in the results of Álvarez-Carretero et al. (2021). In simple terms, this is because in retrospect, lineages reflect the routes to modern day survivors; and clearly, a species living just after a mass extinction is more likely to lead to modern day survivors than one living just before.

**Fig 8.**
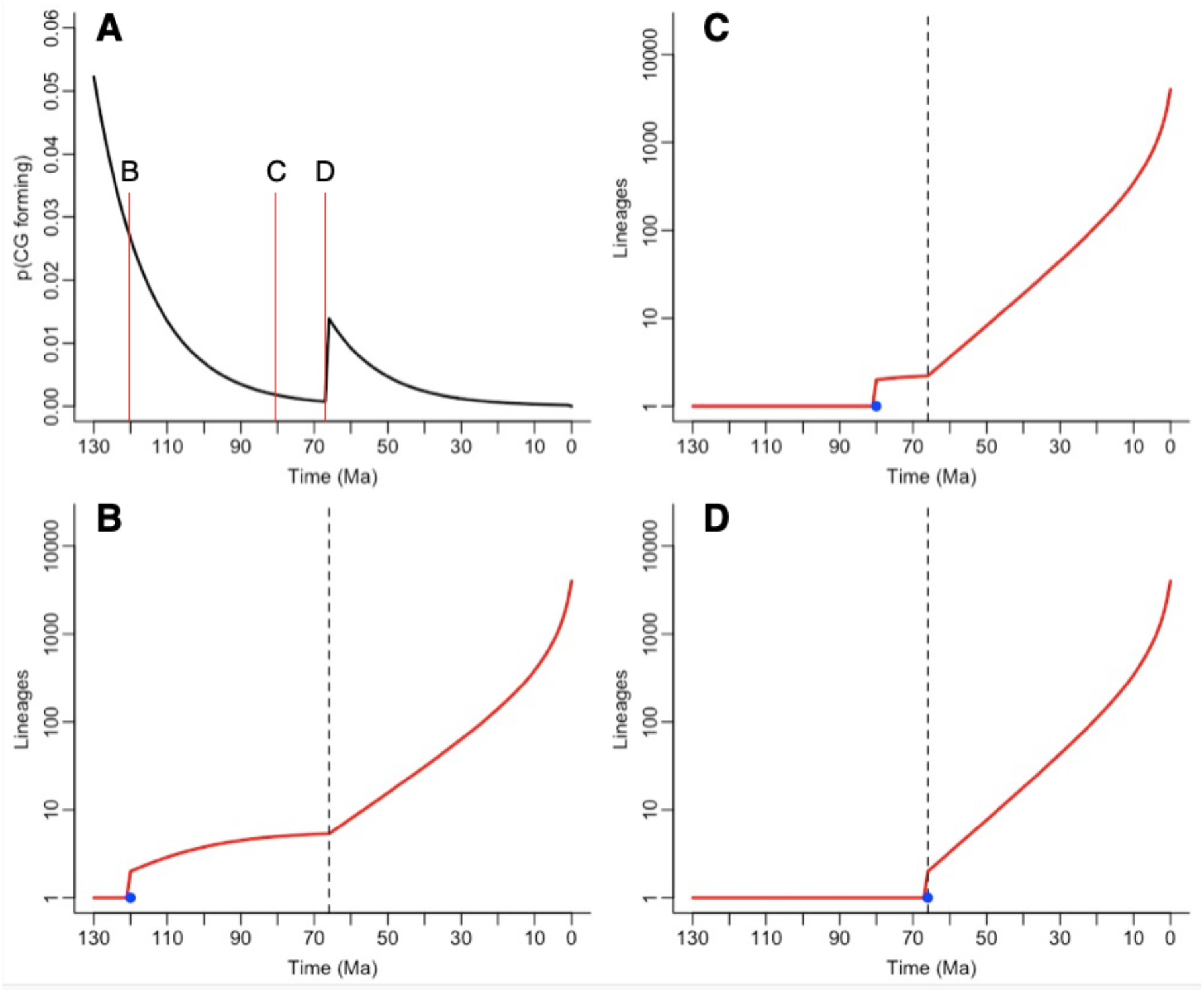
Birth-death modelling through mass extinctions. Here, a clade commencing at c. 130 Ma, and diversifying to give rise to c. 4000 living species with a background extinction rate of 0.5/Myrs passes through a mass extinction of 95% at 66 Ma. **A** The probability of the crown group forming through time. The red lines marked B, C, D indicate the positions of the crown group emergence illustrated in the other three panels. **B** The lineage through time (LTT) plot when the crown group emerges at c. 125 Ma (the blue dot indicates the oldest possible crown group fossil age). **C** The same, but for an emergence at the focal time of Álvarez-Carretero et al. (2021). **D** The same, but for crown group emergence just after the mass extinction. Note that in each case the LTT plot has a distinct inflection at the point of extinction, reflecting the difficulty of coalescence back through a mass extinction.

As well as exploring the LTT plot morphology, it is also possible to produce heat maps using the method of Budd and Mann (2020)^51^ (Fig. 9) for stem- and crown-group diversity through time for each of the scenarios of Fig. 8. These reveal a further difficulty in reconciling the estimates of molecular clocks with the established fossil record. As noted above, there are no uncontroversial crown group placentals older than 66 Ma, but there are thousands of uncontroversial stem group placental fossils before the K-Pg boundary. As shown in Fig. 9, if the crown group of placentals emerged very early, or at c. 80 Ma, we would in both these cases expect a substantial diversification of the crown group before the K-Pg boundary, comparable to the stem diversity at the same time. Given the preservation of *stem-group* placentals during this interval, it is natural to ask where the crown group fossils are^44^. Conversely, if the crown group forms immediately after the K-Pg mass extinction, the expected abundance of stem and crown groups through time closely matches the known distribution of fossils.

**Fig. 9.**
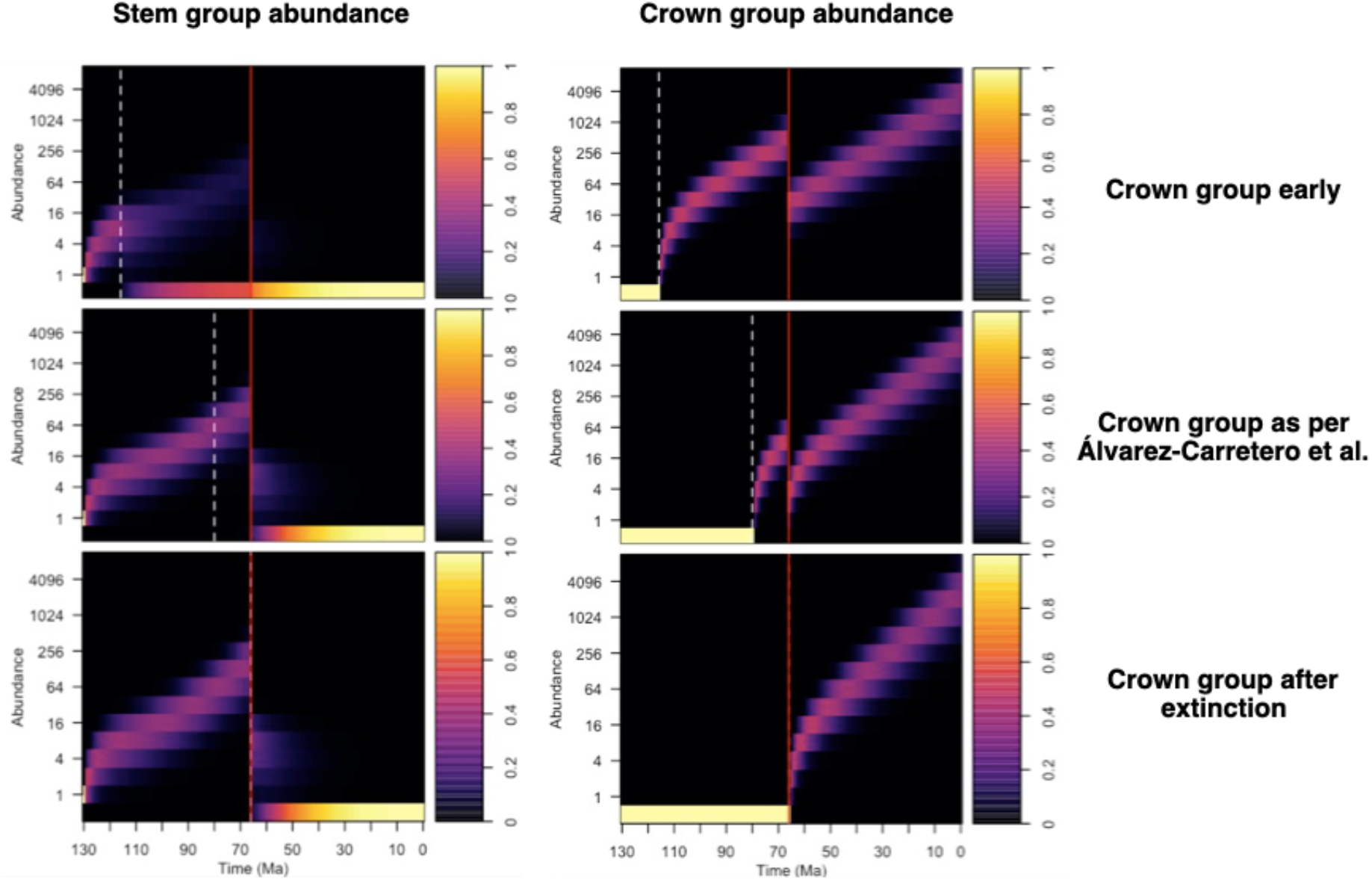
Heat maps for placental diversification for each of the three scenarios in Fig. 8. White dashed line: time of crown-group origin; red dashed line, mass extinction. Note that in both the upper and middle scenarios, significant crown-group diversity should be expected in the Cretaceous, with, in the focal analysis of Álvarez-Carretero et al. (2021), approximately one third of total late-Cretaceous diversity being crown group, and with a total of several hundred species being expected during this interval of time. Conversely, the bottom scenario matches the known fossil record well.

### Conditional calibrations and prior beliefs

Both the fossil record and the living organisms that make up a clade are the result of the evolutionary process of diversification, and it is possible to model this process using birth-death models^54^. The problematic aspect of this modelling is inserting its influence into the priors of Bayesian relaxed clock inference. For example, so-called “multiplicative” priors, implemented in BEAST, use a tree prior based on a birth-death model, but then multiply the priors for nodes that have fossil calibrations with fossil calibration ranges; and this creates improper priors^55^. Conversely, using a fixed tree topology, MCMCTree implements a “conditional” prior, which is based on fossil calibration ranges where they exist; a generic birth-death model is used only for uncalibrated nodes. The conditional prior is formed by truncating the fossil calibration ranges so as to ensure the (assumed) known phylogenetic topology (e.g. that the origin of Bilateria must post-date the origin of Metazoa). This is done by first sampling from each calibration range independently, and then accepting the joint sample over all nodes if those topological constraints are satisfied. The effect of this procedure is very pronounced in the deepest nodes; sampled values for the origin of Metazoa that are anywhere close to the age of the earliest fossil are very likely to be rejected, since it is very likely that the accompanying sample from the Bilateria node (or other deep nodes such as Protostomia) will be older, violating the topological constraints. Conversely, very old samples from the Metazoa range are very likely to be accepted – any sample older than the Lantian is essentially guaranteed to be older than the accompanying sample from Bilateria and other nodes. This skews the resulting prior distribution of all deep nodes to be towards the oldest limit of their calibration range, as they recursively provide more ‘room’ for the nodes above them.

We contend that this procedure produces a prior that does not represent the actual uncertainty we have over the possible node ages. Instead, if one truly believes that Metazoa could, with equal plausibility, have originated anywhere after 833Ma and before the earliest fossil, we think it would be more reasonable for the prior over the age of the Metazoa (rather than the calibration range) to reflect this. In other words, we should have a uniform prior over the total age of the clade (the focal object of estimation), rather than placing this on a calibration density that has little statistical or biological meaning.

There are multiple ways that the full prior structure could be determined. Here we propose one very conservative variation upon that implemented in MCMCTree. We take as a starting point a uniform prior on the age of the basal node. Conditioned on the age of the basal node, we construct the rest of the prior using the conditional prior method. This prior can be sampled (and potentially utilised) in MCMCTree as follows: we take a discrete sample of potential metazoan ages uniformly spaced between the oldest fossil and 833 Ma (we used a 10 Myr spacing of samples). For each of these dates we construct a calibration tree file, with fossil calibrations as before, with the sole difference that the metazoan node is tightly constrained to lie within the chosen sample range (e.g. between 830 Ma and 820 Ma). Samples can then be taken from the prior implied by each of these distinct calibrations, using the usual MCMCTree procedure to generate samples from the prior. The samples from each calibration are then aggregated to produce the overall prior, which by construction is now uniform for the Metazoan age.

The result of implementing this procedure is shown in Fig. 10 for the four deep nodes of Metazoa, Bilateria, Protostomia and Deuterostomia, showing the original calibration ranges, the prior constructed as above and for comparison the prior of Dos Reis et al. The metazoan prior is now uniform by construction, and thus does not exclude potential scenarios where the earliest fossil is close in age to the origin of the clade. While older ages are still favoured for the Bilateria, Protostomia and Deuterostomia, this procedure can be seen to produce a much greater concentration of prior probability on younger origin times for each node than the original prior method. In principle, a similar procedure could be used to arrive at the posterior distribution, by once again running MCMCTree to derive a posterior for each calibration file and then aggregating the results, though this is prohibitively time consuming.

**Fig 10.**
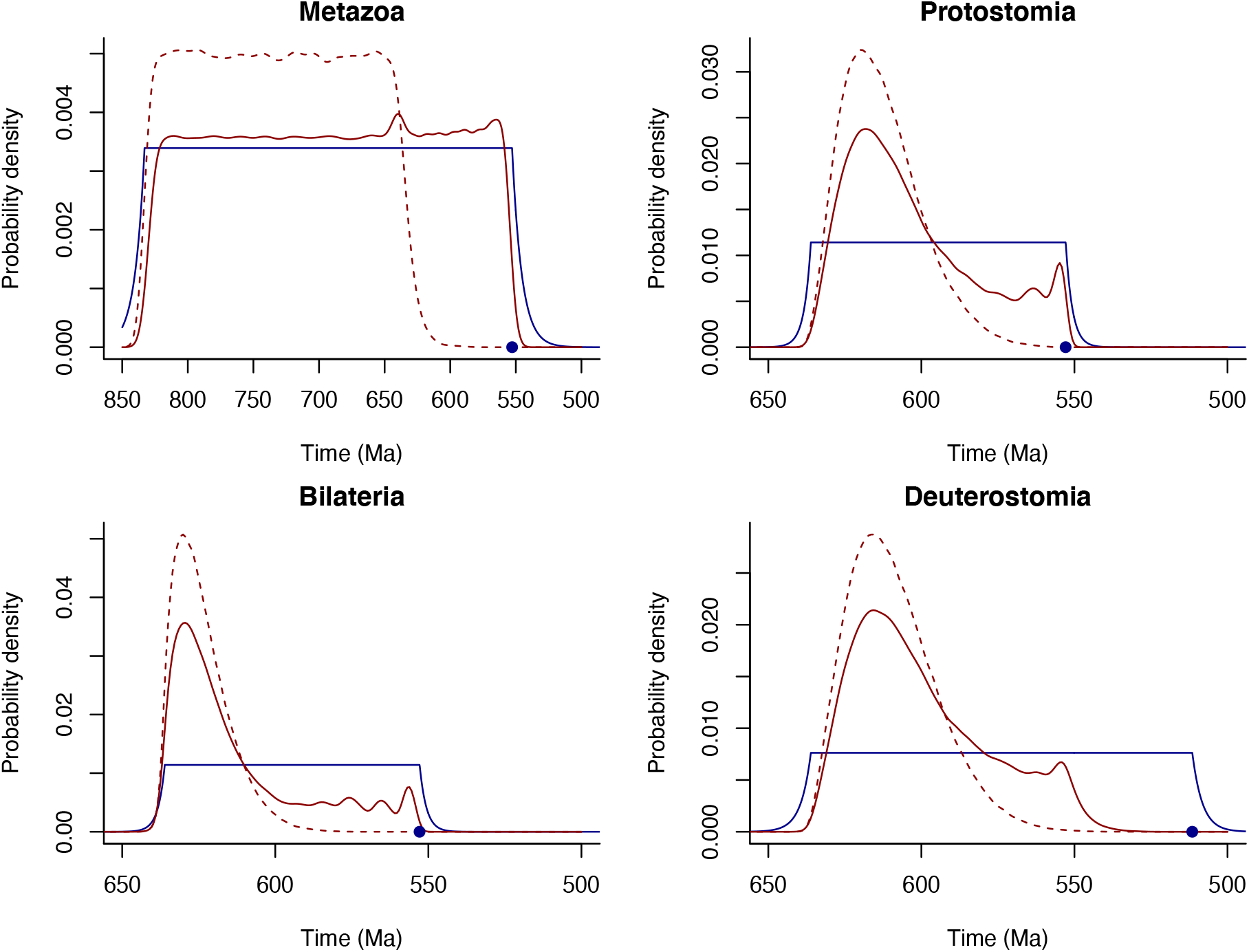
The calibration (blue, oldest fossil blue dot), uniform prior of Dos Reis et al. (dotted red) and the improved prior proposed here (solid red). Note that the solid red prior assigns more probability closer to the age of the oldest fossil.

Although we believe this method of constructing priors from the calibrations is modestly superior to that implemented in MCMCTree (although much more time consuming), it suffers from at least two significant shortcomings. The first is that it still does not have a properly informed model for diversification behind it; and secondly, that it does not take into account what we can learn from the entire fossil record of a clade, not just its oldest fossil. Some attempts at modelling diversification from the fossil record *have* been made^56,57^, such as for primates^57^. This study uses a logistic birth-death model for its model, but it is implemented in a deterministic way which is inappropriate because of the importance of the push of the past^50^. In the later study by Wilkinson et al.^56^, factoring in the K-Pg boundary extinction strongly skews the origin of the placentals to after the boundary.

### Known and unknown unknowns: the philosophy of prior beliefs

As we have previously discussed, we consider the problem of the timing of the origins of clades to be a strongly empirical question. As such, we cannot discover what the “true answer” to when (for example) the metazoans evolved, but rather state what our best hypothesis is based on the available evidence; and this answer will always be corrigible. When assessing this evidence, several levels of uncertainty need to be considered, and these might be categorised as “known” and “unknown” unknowns. A known unknown would, for example, be the age of the oldest member of a clade based on a suitably constructed stratigraphic confidence interval analysis of a known fossil record. Conversely, an “unknown” unknown might be the postulation of as yet hidden intervals of non-preservation in the fossil record, or an interval of evolution when all animals were tiny and impossible to preserve. This distinction is important because only the first type of uncertainty is evidenced, whereas as the second is not. Factoring it in as a possibility into a molecular clock analysis will, however, inevitably enormously increase the mean and range of possible posteriors. We therefore believe that only evidenced uncertainties should be used. This inevitably implies that the correct prior on a node should be its suitably constructed stratigraphic confidence interval unless there is evidence to the contrary.

As we have previously discussed for the case of the metazoans^20^, it is clear that the oldest accepted evidence in the fossil record is that of the Avalonian “Ediacaran” assemblages, aged <575 Ma^23^. Older assemblages such as the Lantian and Weng’an biotas (both probably just older than 600 Ma), although showing exceptional preservation, do not preserve fossils accepted to be animals^20,58,59^. Unless one invokes an “unknown unknown” then, the fossil record clearly indicates that animals evolved shortly before the oldest Ediacaran assemblages, and our results here show that molecular evidence is compatible with this view (by which we mean that if the calibration within MCMCTree reflects this fossil evidence, the molecular analysis does not indicate a contrary date). Similarly, theoretical and empirical considerations both indicate that placental mammals evolved after the K-Pg boundary; and molecular evidence is also compatible with this. We note that if new convincing fossils that are older than this were to be found, or known fossils were convincingly re-evaluated, then these dates would be open to revision. Thus, estimates of ages of clades must always depend primarily on our interpretations of the fossil record, and not on the relatively small influence that relaxed clock methods might have on them.

## Discussion

In this contribution, we have considered two well-known and controversial radiations, that of the animals and the placentals within the mammals. We conclude from our analyses that relaxed molecular clocks do not provide a reliable way of estimating the ages of clades. In both cases, we have shown that the popular MCMCTree methodology using conditional priors on the node ages suffers from a number of well-known and less well-known problems: i) deep nodes tend to have priors with mean ages much older than the raw calibrations would suggest; ii) Such priors assign virtually no probability close to the fossil record, which implies that these clock analyses are not testing the hypothesis that the fossil record might be an accurate reflection of evolutionary events; iii) simply changing the maximum age on the root calibration can have very significant effects on basal node posterior age inference; iv) in general there is a strong dependence of the posterior estimates on the priors. Taken together, the clear implication is that the molecular part of the analysis does not allow us to distinguish between different times of origin of the clade, and thus does not contradict the general picture provided by the fossil record.

An apparent counter-example to this general rule is provided by the placental node in the mammals, because here the posterior *does* strongly differ from the prior; but here the prior can clearly be seen to be in extreme conflict with the molecular data, and is thus exerting an inappropriate effect on the posterior. When a more appropriate prior is used, the prior-posterior relationship, rightly or wrongly, is restored. Because this point might be misunderstood and seen as ‘heads I win, tails you lose’, it is important to clarify: in a sound analysis the priors should be broad enough not to rule out any reasonable hypothesis; the molecular analysis should then refine the prior, leading to greater precision (and ideally accuracy), within the scope of the prior. Posteriors that either fail to refine the prior, or that lie well into the tails of the prior, show that either the molecular evidence is weak or that the construction of the prior is systematically flawed.

The explosive post-Cretaceous model for placental mammal origins has been deprecated, partly on the grounds of the alleged too-rapid rates of molecular evolution that must be inferred for the earliest branches^43^. In addition, in order to fit in the several lineage-production events between the boundary and the first putative diagnosable in-group members of the placentals^43^, several speciation events must have occurred in a few hundreds of thousands of years. If the placentals are a “large clade”^50^ however, then more lineages than expected would be created in the earliest part of a radiation, which in turn implies few plesions. In other words, one can postulate that for such a radiation, the speciations that gave rise to the lineages were a significant proportion of the total number. Even so, it is clear that the short-term speciation rate would be very high under such circumstances, and such rates are not captured by time-homogeneous birth-death models. However, there is no good reason to think that speciation (or extinction) or indeed molecular substitutions are really Poisson distributed^60–62^, and it is entirely plausible that all these rates could be much higher than the long-term mean when short intervals are considered. This should be seen in the context that crown group formation is inherently unlikely shortly before mass extinctions^51^, and that no uncontroversial crown group placental fossils have been found older than 66 Ma. Thus, it is unclear what the grounds really are for thinking that placentals emerged before the boundary; and there are several sound reasons for thinking they emerged after it.

MCMCTree analyses are popular because they can be run on large datasets and are intuitively easy to understand (for example, only a single calibration, rather than mixed birth-death and fossil calibrations is applied to each node). However, this simplicity comes at a price, which is that the calibrations, and in particular the priors that arise from them, are not biologically informed (for example, in general they do not reflect what is known about the dynamics of radiations or the effect of mass extinctions). Here we show that these factors are very important in understanding the timing of origins of clades, and thus should be considered as part of any “total evidence” approach^63^. Dependence of posterior estimates on the priors has been shown for other clades^64^ and other software as well^61,65,66^, suggesting that the issues we raise here are widespread^40^. We suggest that, at the minimum, prior-posterior plots should be included in molecular clock analyses as a matter of routine, so that the total contribution of the calibrations, priors and molecular data can be readily seen.

As noted above, we regard the age of a clade as an empirical matter, and thus any estimate should and must be open to refinement or even refutation from evidence. Furthermore, we regard the fossil record in itself as providing evidence for such timings, albeit of varying quality. We would thus dissent, for example, from the recently-expressed view about angiosperm origins that because molecular clock estimates are not reliable, we thus have no real idea about when they emerged^65^. The ingrained idea that the fossil record can only start after a long and unpredictable “lag” period or “phylogenetic fuse”^67–69^ has been criticised on various grounds^51^, but seems to be the basis for the relative insouciance with which large gaps between clocks and fossils are accepted. When the fossil record is of high quality, with large numbers of fossils from geographically widespread localities, as is the case for the clades we are considering here, then to posit an origin much before the fossil record commences would require the addition of more empirical evidence. We have shown here that, at least in the cases we consider, this evidence cannot come from molecular clock analysis, and it does not come from the fossil record *per se*. Such evidence might include, for example, phylogenetic reconstruction that suggests early members of a clade were all fragile or otherwise hard to preserve. In the absence of clear evidence of this sort, though, the positing of large gaps in the fossil record, especially when they are well outside those suggested by stratigraphic confidence intervals^70,71^, appears to be an unnecessary hypothesis that requires inevitable appeals to *ad hoc* explanations, which although common in the literature, cannot be general phenomena. In this light, molecular clock analyses should in our view attempt to test the fossil-record derived date hypothesis; and they cannot do this if they assign vanishingly small prior probabilities to the eventuality that the fossil record is broadly correct^39^; nor are they useful if they are compatible with both deep and shallow origins of clades, as we show here for the animals and placentals.

## Methods

For the MCMCTree molecular clock analyses, we reproduced the analyses of ref. ^72^ and ref.^19^ as far as possible, using PAML4.8 and PAML4.9 respectively, and the datasets associated with each study. To produce Fig. 9, we used the methods and code of ref. ^51^, setting the background extinction rate to be 0.5/Myr^73^ (speciation rate was set so as to produce an expected 4000 species in the present, commensurate in scale with the number of known placental species).

## Acknowledgements

We thank Richard Copley for some initial help with computing, and Mario Dos Reis Barros for kindly making some material from ref. 16 available.

